# Quantifying Epigenetic Stability with Minimum Action Paths

**DOI:** 10.1101/2020.02.25.964726

**Authors:** Amogh Sood, Bin Zhang

**Affiliations:** Departments of Chemistry, Massachusetts Institute of Technology, Cambridge, MA, USA

## Abstract

Chromatin can adopt multiple stable, heritable states with distinct histone modifications and varying levels of gene expression. Insight on the stability and maintenance of such epigenetic states can be gained by mathematical modeling of stochastic reaction networks for histone modifications. Analytical results for the kinetic networks are particularly valuable. Compared to computationally demanding numerical simulations, they often are more convenient at evaluating the robustness of conclusions with respect to model parameters. In this communication, we developed a second-quantization based approach that can be used to analyze discrete stochastic models with a fixed, finite number of particles using a representation of the *SU* (2) algebra. We applied the approach to a kinetic model of chromatin states that captures the feedback between nucleosomes and the enzymes conferring histone modifications. Using a path integral expression for the transition probability, we computed the epigenetic landscape that helps to identify the emergence of bistability and the most probable path connecting the two steady states. We anticipate the generalizability of the approach will make it useful for studying more complicated models that couple epigenetic modifications with transcription factors and chromatin structure.

---

A remarkable achievement of multicellular organisms is the formation of distinct cell types with identical genomes. Covalent modifications of histone proteins, of which DNA wraps around to form chromatin, are expected to be crucial for the emergence of cellular diversity.^1^ These epigenetic marks can regulate the output of the genome by promoting or restricting the accessibility of the DNA sequence. They are known to impact the openness of chromatin and global genome organization, though the molecular mechanisms are only beginning to emerge.^2–4^ Therefore, multistability in chromatin states formed by various histone modifications or combinations thereof can potentially give rise to distinct patterns of gene expression and inheritable phenotypes.^5–8^ Evidence for bistable and inheritable epigenetic marks has indeed been found that can be attributed to the presence of positive feedback loops wherein nucleosomes that carry a particular modification recruit, either directly or indirectly, enzymes that catalyze the same modification on neighboring nucleosomes.^9–13^

Mathematical modeling of the reaction networks of histone modifications can help determine factors that are crucial for epigenetic stability. Distilling the essence of feedback mechanisms, Dodd and coworkers introduced a simplified kinetic model with bistable chromatin states.^14^ One envisions a system of *N* nucleosomes, where a nucleosome can exist in either a modified or an unmodified state (see Fig. 1). As a first approximation, spatial organization of chromatin is neglected, and the kinetics of the system can be described with the nonlinear dynamics given below

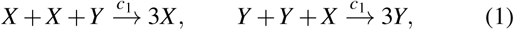

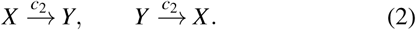

**FIG. 1.**
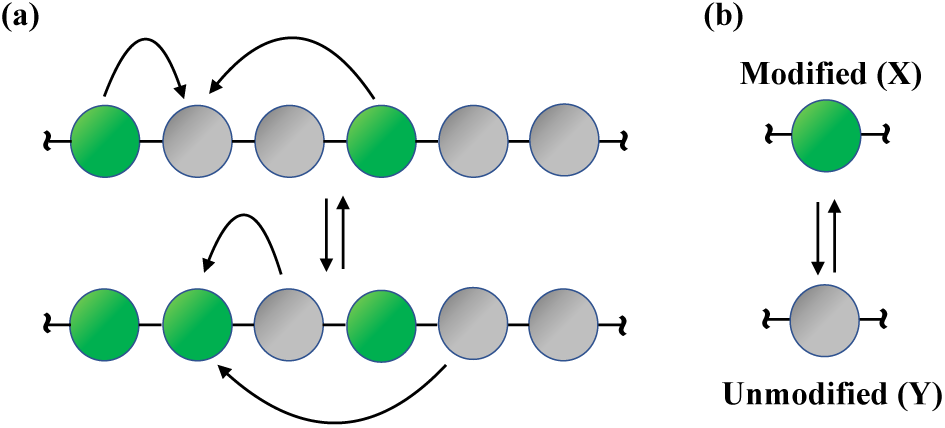
Illustration of the kinetic model for the interconversion between modified (green, *X*) and unmodified nucleosomes (grey, *Y*). (a) Recruited conversion defined in Eq. 1 that requires a pair of (un-)modified nucleosomes to alter the state of a nucleosome. (b) Noisy conversion (Eq. 2) with first order kinetics.

The reactions governing the dynamics in Eq. (1) and (2) represent recruited and thermalized, noisy conversions respectively. From analysis of regulatory circuits, we know that bistability requires not only positive feedback but also nonlinearity in the feedback loop.^9,15–17^ This is encapsulated by the fact that in this model the recruited conversion of *Y* to *X* (or *X* to *Y*) is bimolecular in *X* (or *Y*) and unimolecular in *Y* (or *X*). Thus the rate of production for a given nucleosome type responds to increases in its own concentration in a nonlinear fashion.

Far from being a trifle toy-model, the kinetic scheme above is not unlike the mating type silencing in *S. Cerevisiae*.^14,18–21^ Generalizing the above model to more than 2 epigenetic states has been attempted as well.^21–25^ Their elegance and biological relevance have inspired numerous theoretical studies of these models.^5^ A popular approach used in these studies to investigate epigenetic stability is to posit deterministic rate equations followed by bifurcation analysis. Insight into the switching among chromatin states is missed in such deterministic analyses, however. To study the rare transition events between steady states, Dodd and coworkers introduced an approximate Fokker-Planck equation, from which an epigenetic landscape can be constructed.

In this communication, we present an alternative way to analyze such zero-dimensional models. We turn to the original master equation, which is an exact stochastic description of the underlying process describing the temporal evolution of the system’s configurational probability, and reformulate it using second-quantization (or Fock-Space) methods (Doi-Peliti approach).^26–29^ While canonical approaches rely on *bosonic* creation and annihilation operators, we employ operators that are a representation of the *SU* (2) algebra, in order to treat the constraint that fixes the total number of nucleosome types (*X* + *Y* ≡ *N*) in a more mathematically natural fashion.

The Doi-Peliti method has been successfully employed in the study of reaction-diffusion processes,^30^ gene switches,^31,32^ and other systems.^33^ We outline the main results here and the detailed derivations are consigned to the Supporting Information (SI). The Doi-Peliti approach allows us to reformulate the time evolutiodn of the original master equation as an *imaginary* time Schrödinger equation

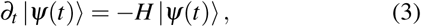

where we have introduced *formally* a state vector |*Ψ*(*t*)⟩ as a superposition of all possible occupation number configurations weighted with their corresponding probabilities (a *generating* function),

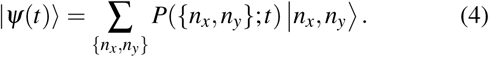

Following standard procedures,^31–33^ *H* is usually expressed in a second-quantized form

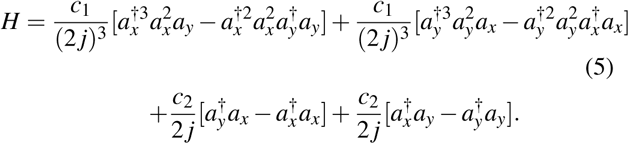

*a*_*i*_ and 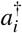 for *i* ∈ {*x, y*} are bosonic creation and annihilation operators that obey the canonical commutation relations

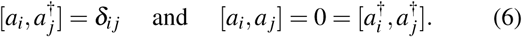

The action of 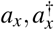 on ket vectors |*n*_*x*_, *n*_*y*_⟩ is given by *a*_*x*_ |*n*_*x*_, *n*_*y*_⟩ = *n*_*x*_ |*n*_*x*_ − 1, *ny*⟩, 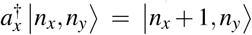, and 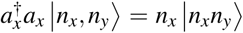. Similar operations can be defined for 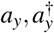. Since for a vacuum state |0, 0⟩, *a*_*i*_ |0, 0⟩= 0, one can obtain any arbitrary ket state as 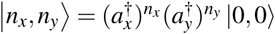.

While it is straightforward to apply the standard formalism up until this point, one notices that the total number of nucleosomes in our system 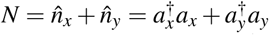 is constant. This is evident from the fact that our Hamiltonian commutes with the total number operator,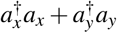. One might also find it slightly philosophically troubling, to use bosonic ladder operators to describe a system with a large, but still finite number of particles. Moreover, the states *X* and *Y* are mutually exclusive, and in conventional quantum mechanics bosonic ladders allow for neither an exclusion principle, nor a cap on the total particle number. However, since the Hamiltonian conserves particle number, we aren’t remiss in our formalism. We can take any equilibrium solution of the master equation and project down to the subspace where the conserved quantity takes a fixed value and we will get another equilibrium solution. In probability theory we would say we are conditioning on the conserved quantity taking a definite value. Thus if our initial state |*Ψ*(0)⟩ is a configuration such that 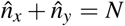 holds true, and then given our prescription of stochastic Hamiltonian all subsequent states will meet this condition as well. However, given that there is essentially only one independent variable, namely *n*_*x*_, we can perhaps phrase this problem in a more natural framework. We develop this framework in the sections that follow.

The starting point of this reformulation is the Jordan-Schwinger map,^34,35^ where we introduce,

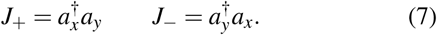

For notational convenience we set *N* = 2 *j*. Here the operators satisfy the commutation relations of *SU* (2) algebra

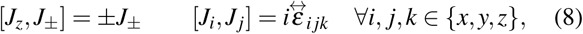

where the structure constant 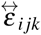 is the Levi-Civita symbol. Re-defining the ket |*n*_*x*_, *n*_*y*_⟩ = |*n*_*x*_, 2 *j n*_*x*_ ⟩ =|*n*⟩, their action is given by,

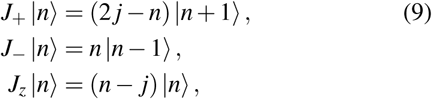

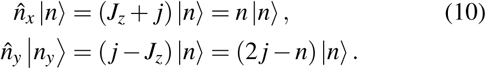

The Hamiltonian can be reformulated as

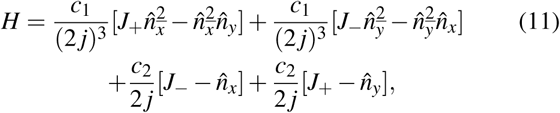

where we have used 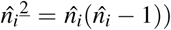 to denote the falling factorial to write down the Hamiltonian in a more compact form.

A great advantage of the second quantization approach is its relative convenience for deriving analytical solutions. For example, a formal solution to Eq. 3 is given by

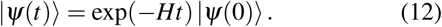

In addition, the transition probability of starting in a state with particle number *n*_*i*_ at time *t* = 0 and ending up in a state with particle number *n*_*f*_ at *t*_*f*_ can be defined as

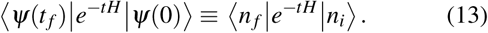

We next seek for a path integral expression of the transition probability that is useful for finding steady states and transition pathways between them. We discretize the time interval [0, *t* _*f*_] into *N*_*t*_ time slices, and then insert a resolution of identity between each time slice. Finally, taking the limit *N*_*t*_ → ∞ we get,

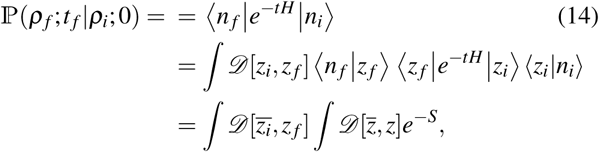

where we have introduced, *ρ* = *n/*2 *j*. By definition, *ρ* represents an order parameter that quantifies the fraction of modified nucleosomes. After performing the integration over 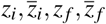. we introduce one final re-parametrization in terms of the density *ρ*. Using

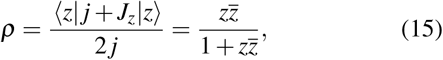

we can rewrite

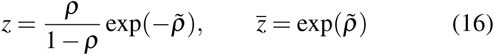

with *ρ*(0) = *ρ*_*i*_ and *ρ*(*t*) = *ρ*_*f*_ and 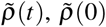 unconstrained. Making these substitutions, the action finally reads,

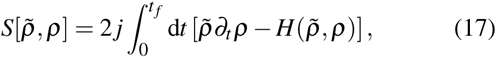

where

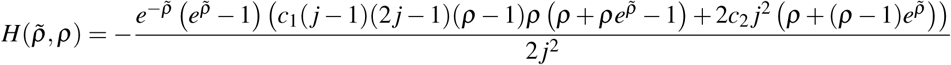

Correspondingly, the time-dependent transition probability (propagator) can be expressed as

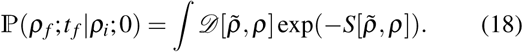

Eq. 17 is the main result of this paper. It allows the computation of both steady state and kinetic results for the model in terms of the order parameter *ρ*. In particular, for 2 *j* ≫ 1, i.e., the small noise regime with many nucleosomes, the path integral in Eq. 18 will be dominated by contributions from the minimum action path.^36–40^ The variational derivatives that minimize the action yield the *classical Hamiltonian equations*

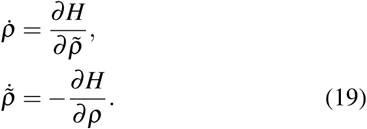

We note that 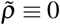 is always a solution to the above Hamiltonian equation. As shown in Figure 2 in blue color, these paths correspond to deterministic dynamics flowing towards steady-state solutions (green dots). The resulting deterministic equation (see Eq. S28 from SI) is identical to that presented in Ref. 14, which was obtained using phenomenological arguments. For the Hamiltonian paths with non-zeros 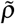, the zero-energy path with *H* ≡ 0 is of particular interest as it represents fluctuations away from the steady states. In bistable regimes, the zero-energy path connects the two steady states via the saddle point (see Fig. 2b red), and corresponds to the maximum likelihood transition path.

**FIG. 2.**
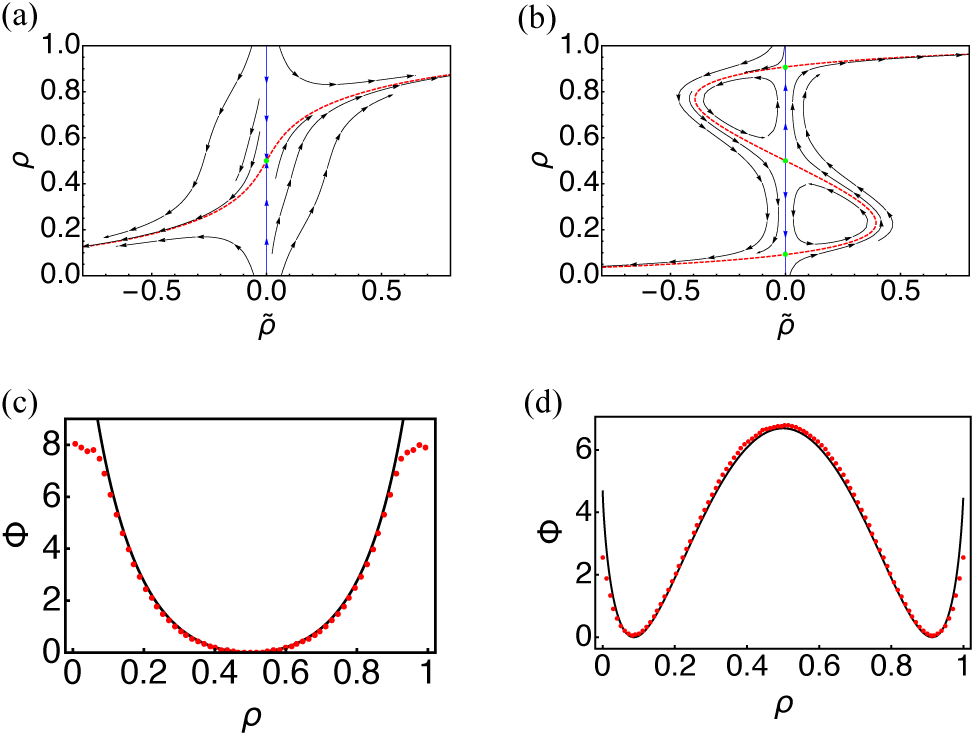
Semi-classical treatment of the path integral expression for transition probability leads to the definition of Hamiltonian equations and quasi-potential. (a,b) Phase portrait determined using Eq. (19) with kinetic parameters *c*_1_*/c*_2_ = 3 (a) and 12 (b). The red dashed lines are zero-energy paths and green dots are steady state solutions. The blue paths represent deterministic trajectories. (c,d) The Friedlin-wentzel quasi-potential (black solid line) compared with the results from stochastic simulations (red dots) for *c*_1_*/c*_2_ = 3 (c) and 12 (d).

**FIG. 3.**
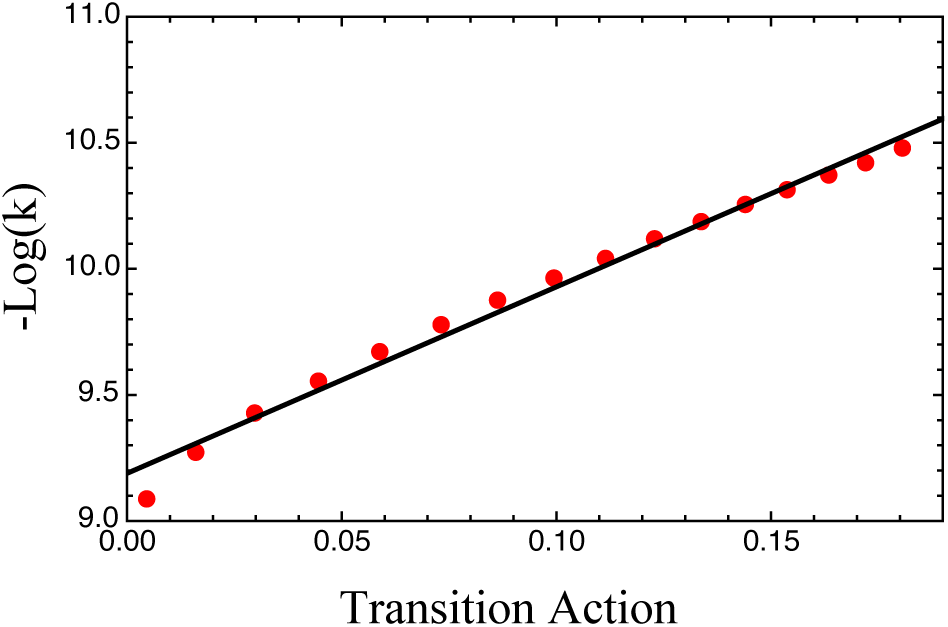
Correlation between the transition rates calculated from the rate equation and from the barrier height of the quasi-potentials for parameter values between *c*_1_*/c*_2_ = 5 and 20.

Quantitative results of the kinetic model can be obtained with the definition of a quasi-potential, F, in the Friedlin-Wentzel sense^41^ in terms of the least-action path (denoting 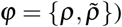),

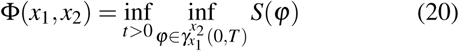

where 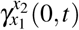 is the set of continuous curves *f* connecting two points *x*_1_, *x*_2_ in configuration space, such that *f* (0) = *x*_1_, *f* (*t*) = *x*_2_. The minima of the Friedlin-Wentzel quasi-potential correspond to attractors of the dynamical system, and the height of the barrier corresponds to the ease of transition between two stable fixed points. As shown in Fig. 2, the quasi-potential correctly captures the emergence of bistability as the parameter *c*_1_*/c*_2_ varies from 3 to 12. *c*_1_ and *c*_2_ are the rate coefficients for recruited and random nucleosome conversions defined in Eqs. 1 and 2. In addition, we found that in both cases, the quasi-potential agrees well with the negative-log of the steady state probability distribution obtained from stochastic simulations (see SI for details). When compared to the rate obtained by the exact diagonalization of the transition matrix of the master equation, the barrier of the saddle point determined from the quasi-potential agrees well with the numerical values over a wide range of parameters. Hamilton’s equation of motion and the quasi-potential, much like their counterparts in classical mechanics, provide intuition regarding the dynamics and landscape of the reaction network.

In this communication, we applied the Doi-Peliti approach to a reaction network that captures the emergence of epigenetic stability from histone modifications.^14^ Together with a transformation enabled by the *SU* (2) algebra, it allowed for the derivation of analytical results that rigorously account for the constraints imposed by a fixed number of particles. The semi-classical treatment of the path integral expression for transition probability further provided a fresh view of the stochastic reaction network in the guise of a “pseudo-mechanical system”. The presented theoretical framework is powerful and general. Its application to more complicated cases with coupled reaction networks, for example, a chromatin switch coupled to a self-activating gene, is underway.

## I. SUPPLEMENTARY MATERIAL

See supplementary material for additional details regarding analytical derivations and numerical simulations.

## II. ACKNOWLEDGEMENT

This work was supported by the National Institutes of Health (Grant 1R35GM133580-01). A.S. acknowledges financial support from the Lester Wolfe and George H Büchi/Kin-Chun T. Luk fellowships.

The data that support the findings of this study are available from the corresponding author upon reasonable request.

