## Supporting Information for "Quantifying Epigenetic Stability with Minimum Action Paths"

### I. CONSTRUCTING THE COHERENT STATES PATH INTEGRAL

For completeness let us start with the master equation for our system (with $N = 2j$),

$$\begin{aligned}\partial_t P(n_x,n_y) = &\frac{c_1}{(2j)^3}[(n_x-1)(n_x-2)(n_y+1)P(n_x-1,n_y+1) - n_x n_y (n_x-1)P(n_x,n_y)] \\ &+\frac{c_1}{(2j)^3}[(n_y-1)(n_y-2)(n_x+1)P(n_x+1,n_y-1) - n_x n_y (n_y-1)P(n_x,n_y)] \\ +\frac{c_2}{2j}[(n_x+1)P(n_x+1,n_y-1) - n_x P(n_x,n_y)] &+ \frac{c_2}{2j}[(n_y+1)P(n_x-1,n_y+1) - n_y P(n_x,n_y)].\end{aligned} \tag{S1}$$

As mentioned in the main text, following the second quantization approach[1–5] and by introducing the state vector $|\psi(t)\rangle$, the time evolution of the original master equation can be recast into an *imaginary* time Schrödinger equation

$$\partial_t |\psi(t)\rangle = -H|\psi(t)\rangle. \tag{S2}$$

The Hamiltonian is defined in terms of the bosonic creation and annihilation operators as

$$\begin{aligned}H = &\frac{c_1}{(2j)^3}[a_x^{\dagger 3} a_x^2 a_y - a_x^{\dagger 2} a_x^2 a_y^\dagger a_y] + \frac{c_1}{(2j)^3}[a_y{}^{\dagger 3} a_y^2 a_x - a_y^{\dagger 2} a_y^2 a_x^\dagger a_x] \\ &+\frac{c_2}{2j}[a_y^\dagger a_x - a_x^\dagger a_x] + \frac{c_2}{2j}[a_x^\dagger a_y - a_y^\dagger a_y].\end{aligned} \tag{S3}$$

Details of the bosonic operators can be found in the main text.

Now we can begin reformulating this system using the Jordan-Schwinger map.[6] For convenience, let us define the following vectors

$$\boldsymbol{a} \equiv \begin{pmatrix} a_x \\ a_y \end{pmatrix}, \qquad \boldsymbol{a}^\dagger \equiv \begin{pmatrix} a_x^\dagger \\ a_y^\dagger \end{pmatrix}. \tag{S4}$$

Then applying the Jordan transformation one gets,

$$\begin{aligned} Q_\mu &= \boldsymbol{a}^\dagger \boldsymbol{\sigma}_\mu \boldsymbol{a} \\ J_\pm &= Q_1 \pm i Q_2 \\ J_+ = a_x^\dagger a_y \qquad J_- &= a_y^\dagger a_x, \end{aligned} \tag{S5}$$

where $\sigma_\mu$ denotes the usual Pauli Matrices. Also for convenience, we define the following auxiliary operators,

$$J_x = \frac{(J_+ + J_-)}{2} \qquad J_y = \frac{(J_+ - J_-)}{2i} \qquad [J_+, J_-] = 2J_z. \tag{S6}$$

The operators $J_\pm$ satsify the same commutation relations as $SU(2)$ algebra, namely

$$[J_z, J_\pm] = \pm J_\pm \qquad [J_i, J_j] = i\overleftrightarrow{\varepsilon}_{ijk} \quad \forall i,j,k \in x,y,z \tag{S7}$$

We now have have $(2j+1)$-dimensional representation of the $SU(2)$ group. The usual magnetic quantum number is now related to the occupation number $n_x \equiv n$, via $m = n_x - j$. Re-defining the ket $\left|n_x, n_y\right\rangle = |n_x, N-n_x\rangle \equiv |n\rangle$, we have

$$J_+|n\rangle = (2j-n)|n+1\rangle \qquad J_-|n\rangle = n_x|n-1\rangle \qquad J_z|n\rangle = (n-j)|n\rangle \tag{S8}$$

$$\hat{n}_x|n\rangle = (J_z+j)|n\rangle = n|n\rangle \qquad \hat{n}_y\left|n_y\right\rangle = (j-J_z)|n\rangle = (2j-n)|n\rangle\,. \tag{S9}$$

Using these rules, the stochastic pseudo-hamiltonian can now be reformulated as

$$\begin{aligned} H = \frac{c_1}{(2j)^3}[J_+\hat{n}_x^2 - \hat{n}_x^2\hat{n}_y] + \frac{c_1}{(2j)^3}[J_-\hat{n}_y^2 - \hat{n}_y^2\hat{n}_x] \\ + \frac{c_2}{2j}[J_- - \hat{n}_x] + \frac{c_2}{2j}[J_+ - \hat{n}_y]. \end{aligned} \tag{S10}$$

In order to now construct a path integral for the transition probability, one introduces the following left and right spin-coherent states,[7,8] and a resolution of identity,

$$\begin{aligned} |z\rangle &= \frac{1}{(1+z\bar{z})^j} e^{zJ_+}|0\rangle = (1+z\bar{z})^{-j}\sum_{0\le n}^{2j}\binom{2j}{n} z^n|n\rangle\,, \\ \langle z| &= \frac{1}{(1+z\bar{z})^j}\langle 0| e^{\bar{z}J_-} = (1+z\bar{z})^{-j}\sum_{0\le n}^{2j}\langle n|\bar{z}^n, \end{aligned} \tag{S11}$$

$$\int \frac{2j+1}{\pi}\frac{d^2z}{(1+z\bar{z})^2}|z\rangle\langle z| = \mathbb{1}. \tag{S12}$$

The details and subtleties regarding the construction of spin coherent state path integrals have been discussed in the literature.[9–13] However, for completeness we give a brief overview. Starting with the propagator between two normalized coherent states, $\left\langle z_f\middle|e^{-tH}\middle|z_i\right\rangle$, we discretize the time interval $[0,t]$ into $N_t$ time slices, and then insert a resolution of identity of the form (S12) between each time slice. Finally, taking the limit $N_t \to \infty$ we get,

$$\left\langle z_f\middle|e^{-tH}\middle|z_i\right\rangle = \int \mathscr{D}[\bar{z},z]\exp(-S) \tag{S13}$$

where,

$$S = -j\log\frac{(1+\bar{z}_f z(t))(1+\bar{z}(0)z_i)}{(1+\bar{z}_f z_f)(1+\bar{z}_i z_i)} + 2j\int_0^t \mathrm{d}t\left[\frac{1}{2}\frac{\bar{z}\dot{z} - \dot{\bar{z}}z}{1+\bar{z}z} - H(\bar{z},z)\right], \tag{S14}$$

and $H(\bar{z},z) = \langle z|H|z\rangle$. Next, we derive an expression for the *physical* propagator between $\langle n_f|$ and $|n_i\rangle$ representing states of fixed initial and final number of particles respectively. This represents the probability of starting in state with particle number $n_i$ at time $t=0$ and ending up in a state with particle number $n_f$ at $t_f$. To do so one takes,

$$\mathbb{P}(n_f;t_f|n_i;0) = \langle n_f|e^{-tH}|n_i\rangle = \int \mathscr{D}[z_i,z_f] \langle n_f|z_f\rangle \langle z_f|e^{-tH}|z_i\rangle \langle z_i|n_i\rangle . \tag{S15}$$

To get the *physical* propagator from (S13) one needs to subtract $\log\langle z_i|n_i\rangle + \log\langle n_f|z_f\rangle$ from the action, and then integrate over $z_i, \bar{z}_i, z_f, \bar{z}_f$. Then,

$$\mathbb{P}(\rho_f;t_f|\rho_i;0) = \int \mathscr{D}[\bar{z_i},z_f] \int \mathscr{D}[\bar{z},z] e^{-S}, \tag{S16}$$

where we have introduced, $\rho = n/2j$. Here $S$ is now given by,

$$\begin{aligned} S = &-j\log\Big[(1+\bar{z}_f z(t)(1+\bar{z}(0)z_i)\Big] + 2j\int_0^t \mathrm{d}t^{t'}\left[\frac{1}{2}\frac{\bar{z}\dot{z}-\dot{\bar{z}}z}{1+\bar{z}z} - H(\bar{z},z)\right] \\ &+2j\Big[-\rho_i\log\bar{z}_i - \rho_F\log\bar{z}_f + \rho_f\log\rho_f + (1-\rho_f)\log\big(1-\rho_f\big) \\ &+\log\big((+\bar{z}_f z_f)(1+z_i\bar{z}_i)\big)\Big]. \end{aligned} \tag{S17}$$

Now we can integrate over $z_i, z_f, \bar{z}_i, \bar{z}_f$ using the saddle-point method. The derivatives of (S17) fix the initial and final conditions,

$$\frac{1}{2j}\frac{\partial S}{\partial z_i} = \frac{\bar{z}_i}{1+\bar{z}_i z_i} - \frac{\bar{z}(0)}{1+\bar{z}(0)z_i}, \tag{S18}$$

$$\frac{1}{2j}\frac{\partial S}{\partial \bar{z}_f} = \frac{z_f}{1+\bar{z}_f z_f} - \frac{z(t)}{1+z(t)z_f}, \tag{S19}$$

$$\frac{1}{2j}\frac{\partial S}{\partial \bar{z}_i} = \frac{z_i}{1+\bar{z}_i z_i} - \frac{\rho_i}{\bar{z}_i}, \tag{S20}$$

$$\frac{1}{2j}\frac{\partial S}{\partial z_f} = \frac{\bar{z}_f}{1+\bar{z}_f z_f} - \frac{\rho_f}{\bar{z}_f}. \tag{S21}$$

Thus, we get the following four conditions $\bar{z}(0) = \bar{z}_i$, $z(t) = z_f$, $\rho_i = \frac{\bar{z}_i z_i}{1+\bar{z}_i z_i}$ and $\rho_f = \frac{\bar{z}_f z_f}{1+\bar{z}_f z_f}$. After performing the integration the action now reads,

$$S = 2j\left[\frac{z\bar{z}}{1+z\bar{z}}\log\bar{z} - \log(1+z\bar{z})\right]\Bigg|_f^i + 2j\int_0^t \mathrm{d}t^{t'}\left[\frac{1}{2}\frac{\bar{z}\dot{z}-\dot{\bar{z}}z}{1+\bar{z}z} - H(\bar{z},z)\right]. \tag{S22}$$

We introduce one final re-parametrisation in terms of the density $\rho$. Using

$$\rho = \frac{\langle z|j+J_z|z\rangle}{2j} = \frac{z\bar{z}}{1+z\bar{z}}, \tag{S23}$$

one rewrites

$$z = \frac{\rho}{1-\rho}\exp(-\tilde{\rho}), \qquad \bar{z} = \exp(\tilde{\rho}) \tag{S24}$$

with $\rho(0) = \rho_i$ and $\rho(t) = \rho_f$ and $\tilde{\rho}(t)$, $\tilde{\rho}(0)$ unconstrained. The Jacobian for the change of variables is $(1-\rho)^2$, and $\int \frac{2j+1}{\pi}\frac{d^2z}{(1+z\bar{z})^2}$ is replaced by $\int \frac{2j+1}{\pi}d^2\rho$. Then,

$$S[\tilde{\rho},\rho] = 2j\int_0^t \mathrm{d}t'\,\tilde{\rho}\partial_t\rho - H(\tilde{\rho},\rho) \tag{S25}$$

where,

$$H(\tilde{\rho},\rho) = -\frac{e^{-\tilde{\rho}}\left(e^{\tilde{\rho}}-1\right)\left(c_1(j-1)(2j-1)(\rho-1)\rho\left(\rho+\rho e^{\tilde{\rho}}-1\right)+2c_2j^2\left(\rho+(\rho-1)e^{\tilde{\rho}}\right)\right)}{2j^2}$$

and the transition probability (propagator) is

$$\mathbb{P}(\rho_f;t_f|\rho_i;0) = \int \mathscr{D}[\tilde{\rho},\rho]\exp(-S[\tilde{\rho},\rho]). \tag{S26}$$

To get the deterministic rate equations, we evaluate,

$$\dot{\rho} = \left.\frac{\partial H}{\partial\tilde{\rho}}\right|_{\tilde{\rho}=0} = \frac{(1-2\rho)\left(c_1(j-1)(2j-1)(\rho-1)\rho+2c_2j^2\right)}{2j^2} \tag{S27}$$

which for $j \gg 1$, gives

$$\dot{\rho} = (1-2\rho)(c_1(\rho-1)\rho+c_2). \tag{S28}$$

This equation is in agreement with Micheelsen *et.al.*,[14] and it yields a single stable real fixed point, $\rho^* = 0.5$ when $c_1/c_2 < 4$ and an unstable fixed point at $\rho^* = 0.5$ and 2 stable fixed points at $\rho^* = 0.5 \pm 0.5\sqrt{(c_1-4c_2)/c_1}$ when $c_1/c_2 > 4$.

### II. DETAILS OF NUMERICAL SIMULATIONS

Stochastic simulations of the chromatin state model with $N = 60$ nucleosomes were carried out using the Gilesspie algorithm.[15] The rate coefficient for random nucleosome conversion defined in Eq. 2 of the main text is set as $c_2 = 1\tau^{-1}$, where $\tau$ is the time unit. The value for $c_1$ was varied to explore the stability of the model as detailed in the main text. For each value of $c_1$, we performed $10^4$ simulations, each of which lasted for $10^5\tau$. These trajectories were initialized with random configurations, and the number of modified nucleosomes along each trajectory was recorded at every $10^2\tau$. We then combined the values from all trajectories to estimate the steady state probability distributions, $P_{ss}$, shown in Figure 2 of the main text.
